# Neuroimaging contrast across the cortical hierarchy is the feature maximally linked to behavior and demographics

**DOI:** 10.1101/730226

**Authors:** Feng Han, Yameng Gu, Gregory L Brown, Xiang Zhang, Xiao Liu

## Abstract

We employed a data-driven canonical correlation analysis to investigate the population covariance of whole-brain cortical thickness, resting-state functional connectivity, and hundreds of behavioral/demographic measures in a large cohort of individuals. We found that the maximal thickness-behavior correlation and the maximal connectivity-behavior correlation are largely converged along the same direction across subjects, which is characterized by very specific modulations of all three modalities. Along this direction, individuals tend to have more positive and less negative behavioral/demographic traits, and more importantly, their functional connectivity and cortical thickness show a similar divergent modulation across the cortical hierarchy: thinner cortex and stronger functional connectivity at the higher-order cognitive regions whereas thicker cortex and weaker connectivity at the lower-order sensory/motor areas. These findings provide a unique link between structural and functional brain organizations and human behavior. Specifically, they suggest that the cross-hierarchy contrast of structural and functional brain measures may be a specific feature linked to the overall goodness of behavior and demographics.

---

Understanding the neural and structural basis of behavioral variability across individuals is an important task for neuroscience. Modern neuroimaging research has made significant progress towards this goal through the study of the brain-behavior relationship across individuals (1–3). The advent of resting-state functional magnetic resonance imaging (fMRI) enabled a non-invasive measurement of functional brain connectivity (4, 5), which has been linked to a variety of individual measures, such as demographic factors (6), lifestyle (7), and cognitive functions (8, 9). A comprehensive study of this connectivity-behavior relationship was recently made possible by the Human Connectome Project (HCP), which collected resting-state fMRI data along with hundreds of behavioral, cognitive, psychometric, and demographic measures from a large cohort of 1,200 subjects (10). A data-driven canonical correlation analysis (CCA) (11) to the HCP data revealed only a single significant CCA mode of population covariation that maximally links individuals’ functional connectivity with their demographic/behavioral measures. Interestingly, a characteristic pattern of behavioral changes emerges in this mode direction with individuals showing more positive subject traits, such as good performance on memory and cognitive tests, high life satisfaction, and good education, towards one end but more negative ones, such as substance use, rule-breaking behavior, and anger, towards the other (12).

However, structural and morphological brain changes along this specific “positive-negative” mode maximally linking the functional connectivity and behavioral/demographic measures remains unknown, even though both modalities have been linked to structural/morphometric brain properties separately. For example, the resting-state connectivity has been linked to structural connectivity measured with diffusion imaging (13), micro-structural properties such as cortical myelination (14), and morphometric measures such as the cortical thickness (15). On the other hand, inter-subject variability in brain structures and morphology, particularly the cortical thickness, has been repeatedly linked to individual differences in various subject measures, including intelligence, aggression, and life satisfaction (3, 16–22). Therefore, it would be interesting to know whether this positive-negative mode of connectivity-behavior covariation is associated with changes of certain structural/morphometric brain properties. If so, would this association also represent the maximal correlation between this structural/morphometric property and the other two modalities? Most importantly, just like a specific cognitive function is often linked to structural and functional features of a specific brain network or region, can we link the overall goodness of subject measures, varying along the “positive-negative” mode, to structural and functional features of any specific brain area or network?

Here we seek the answer to the above questions by studying the population covariation of three distinct modalities of data: cortical thickness (a morphometric brain property that has been repeatedly linked to various behavioral aspects), functional connectivity, and behavioral and demographic measures, in a large cohort of HCP subjects. We simply apply CCA to all pairs of the three modalities and find their maximal correlations are approximately aligned in the same “positive-negative” mode direction. More importantly, the overall negative-to-positive change of subject measures along this mode direction is associated with a similar divergent pattern of modulations in both cortical thickness and functional connectivity: the thinner cortex and increased functional connectivity at the higher-order brain regions in charge of more complex cognitive functions whereas the thicker cortex and reduced functional connectivity at the lower-order sensory/motor areas. These results provide new insight into the three-way relationship among brain structure, brain activity, and human behavior by suggesting that the cross-hierarchy contrast between the lower- and higher-order brain regions is the key feature, at least for both cortical thickness and functional connectivity, closely linked to a specific aspect, i.e., the overall goodness, of human behavioral and demographic measures, and this relationship represents the maximal correlations found among the three modalities of data.

## Results

We used multimodal data from 818 HCP subjects who have completed all four resting-state fMRI sessions. We summarized the cortical thickness over the entire cortex (59,412 vertices), resting-state functional connectivity among 200 distinct brain regions (19,900 connections) parceled with independent component analysis (ICA) (23), and 129 non-imaging behavioral and demographic measures (we refer to as subject measures thereafter) for all the subjects. We then applied CCA, a procedure to seek maximal correlations and corresponding linear combinations of any two sets of variables, to all pairs of the three modalities of data after regressing out potential confounds (including age, gender, head motion, brain size, height, weight, blood glucose level, and blood pressure) and reducing non-subject dimension to 100 using the principal component analysis (PCA) (24). All the procedures above were adapted from the previous study that identified the positive-negative CCA mode between the functional connectivity and behavioral/demographic measures (12).

### The maximal thickness-behavior correlation is associated with characteristic changes of both modalities

The application of CCA to the cortical thickness and behavioral data revealed a single highly significant (*r* = 0.70, *p* < 1.0×10^−5^, corrected for multiple comparisons across all modes estimated, **Fig. 1A** and **1B**) mode that represents the maximal correlation between the two. The canonical weights of behavioral measures, i.e., the correlations between single subject measures and the behavior canonical variate, show a pattern similar to what has been found for the connectivity-behavior CCA mode (12): the subject measures commonly considered as positive traits, e.g., the fluid intelligence and life satisfaction, are positively correlated with the CCA mode whereas the measures of negative traits, e.g., marijuana use and behavioral aggression, tend to show negative correlations (**Fig. 1C**). In fact, the most strongly correlated subject measures, particularly those positive traits, are overlapped with those showing the top correlations with the connectivity-behavior mode, including the vocabulary scores, fluid intelligence, life satisfaction, and tobacco use (12). In summary, the cortical thickness co-varies maximally with the behavioral/demographic measures in a direction also featuring the positive-negative behavioral change.

**Fig. 1.**
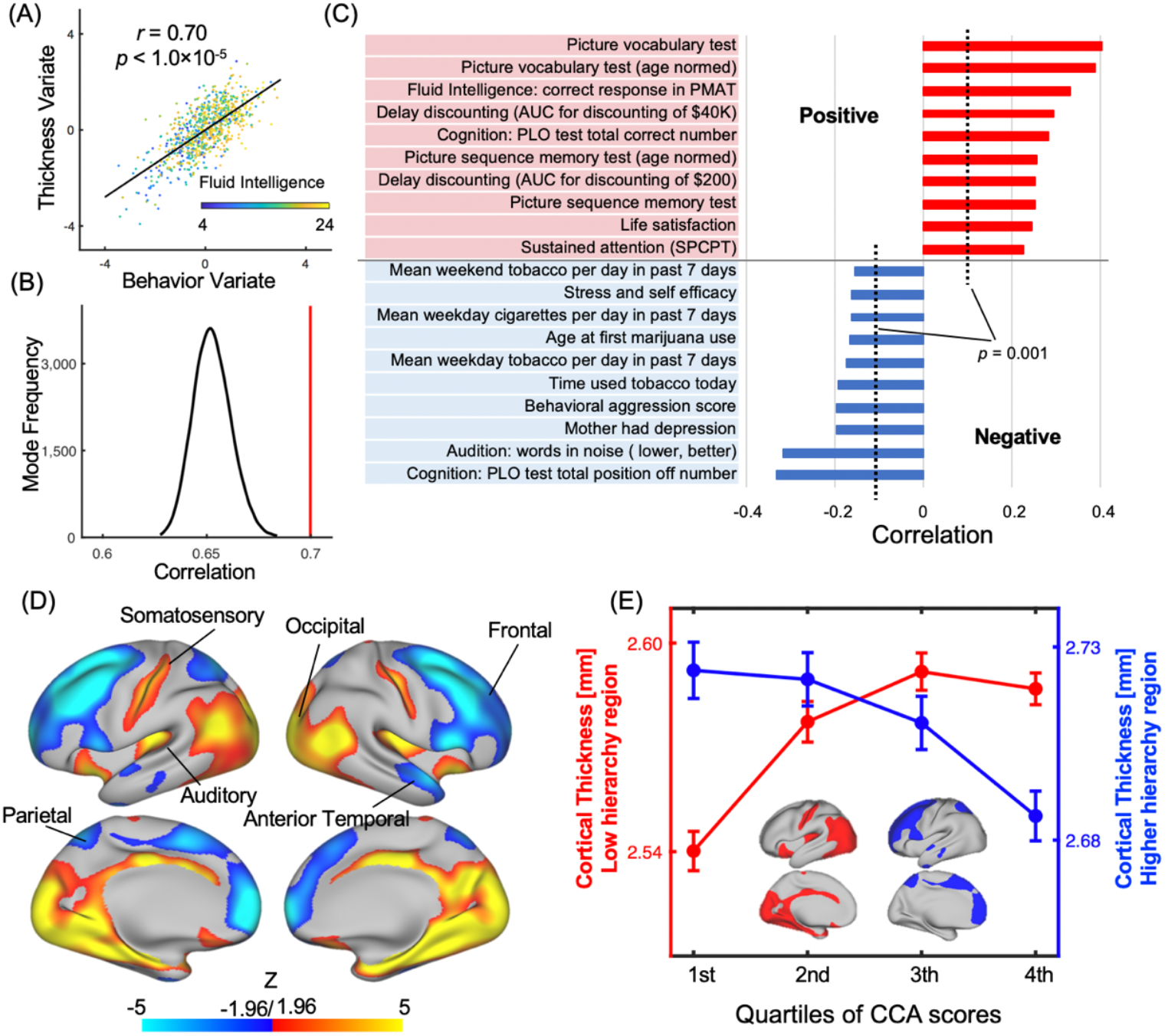
The CCA mode linking the cortical thickness and behavioral/demographic measures. (**A**) A scatter plot shows the correlation (*r* = 0.70, *p* < 1.0×10^−5^) between the two canonical variates of the identified thickness-behavior CCA mode, with one dot representing a subject. Fluid intelligence scores are color-coded the dots of the plot. (**B**) The null distribution of CCA correlations (the most significant pair) between the cortical thickness and 129 subject measures with permuting subject ID 100,000 times for the subject measures. The red line indicates the CCA correlation without permutation, i.e., the identified thickness-behavior CCA mode. (**C**) The top subject measures most strongly correlated with the identified thickness-behavior CCA mode. The positively correlated subject measures (red) generally describe positive personal traits, whereas the negatively correlated subject measures (blue) are commonly perceived as negative personal traits. (**D**) The correlations between local cortical thickness and the identified CCA mode. The original correlations are converted to Fisher’s z scores. Positive correlations (red-yellow colors) are mostly seen at the lower-level sensory/motor regions, whereas negative correlations (blue-cyan colors) are more dominant at the higher-order cognitive brain regions. (**E**) The mean cortical thickness of the lower-order (red) and higher-order (blue) cortical regions in four subgroups of subjects divided by the quartiles of the CCA score, i.e., the mean of the thickness and behavior canonical variates. Error bars represent the standard error of mean (SEM) across subjects.

More interestingly, the canonical weights of the cortical thickness, i.e., the correlations of local cortical thickness (at single vertices) with the identified thickness-behavior CCA mode, revealed a striking spatial pattern of divergent modulations across cortical hierarchy (**Fig. 1D**). Specifically, significant positive correlations are seen mostly at lower-order sensory/motor areas, including the sensorimotor, auditory, and visual cortices, whereas strong negative correlations appear mostly at higher-order brain regions, including the frontal, anterior temporal, and parietal cortices that encompass the most parts of the default mode network (DMN). The contrast is evident across hierarchies in the neocortex, even though positive correlations are also seen at some parts of the allocortex and mesocortex, such as the hippocampal formation and olfactory bulb. In other words, towards one end of this CCA mode individuals tend to show more positive subject traits, thinner cortex at the higher-order cortical regions, but thicker cortex at the lower-order sensory regions (**Fig. 1E**). This thickness-behavior relationship is not driven by any specific subject measures since leaving any one of them out of the analysis produced almost no changes in the final results. Moreover, single subject measures show much weaker correlations with the cortical thickness, which do not show any noticeable pattern of cross-hierarchy divergence (**Fig. S1**). We also conducted a split-half reliability test by randomly dividing subjects into two halves (409 subjects for each) and repeating the CCA analysis on both groups. Similar results, particularly the spatial pattern of cortical thickness weights, were obtained on both groups even with reduced statistical power (**Fig. S2**).

### Cross-hierarchy contrast of the cortical thickness is tightly linked to behavioral measures

The above results imply that the cortical thickness difference between the lower- and higher-order brain regions could be more closely linked to subject measures than region-based statistics. To test this hypothesis, we compared the cross-hierarchy contrast and region-specific cortical thickness in terms for their correlation with behavioral/demographic measures. In addition to the cortical thickness ratio between the lower- and higher-order regions, we calculated the mean cortical thickness of 66 predefined regions (25), of the lower- and higher-order regions, and the cortical thickness ratio between two sets of control brain regions, which were obtained by randomly rotating the masks for the lower- and higher-order brain regions on the brain surface. Compared with the other thickness measures, the cortical thickness ratio between the lower- and higher-order regions showed much stronger correlations with the subject measures (**Fig. 2A** and **2B**). In addition, the correlation strength is consistently higher for this cross-hierarchy thickness ratio than any other thickness measures for any subject measures showing significant correlation to both (**Fig. 2C**). In summary, the cross-hierarchy contrast of cortical thickness appears to be a feature more tightly coupled with the overall subject measures than any region-based measures.

**Fig. 2.**
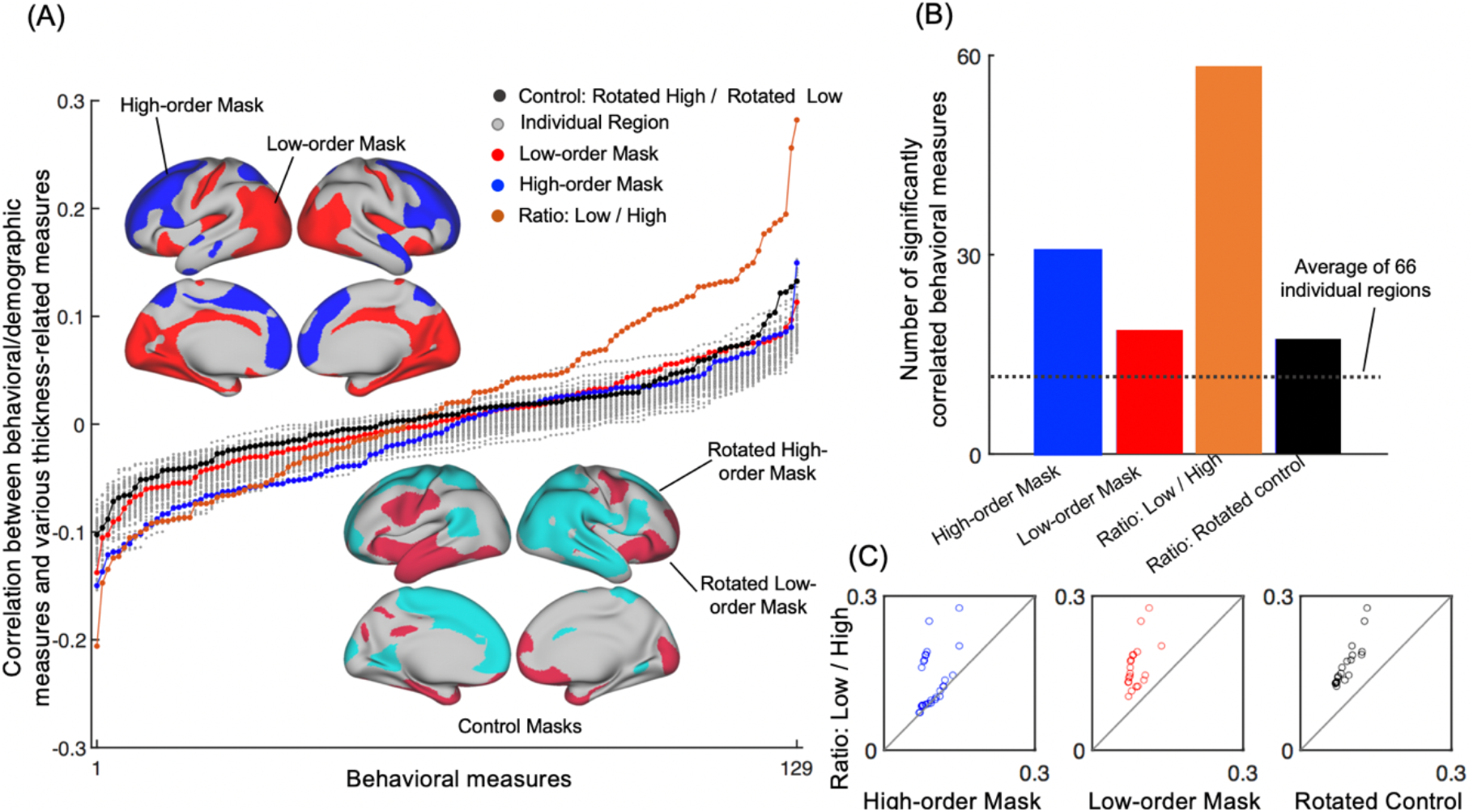
Correlations between behavioral/demographic measures and various thickness measures. (**A**) Sorted correlations between 129 behavioral/demographic measures and 70 different thickness measures, including the mean cortical thickness of 66 pre-defined brain parcels (25) (gray lines), of the lower-(red line) and higher-order (blue line) masks, the thickness ratio between these two masks (orange line), as well as the thickness ratio between two control masks obtained by randomly rotating the lower- and higher-order masks on the brain surface (black line). (**B**) The number of significantly correlated subject measures (*p* < 0.05, uncorrected) was counted for four thickness measures. The thickness ratio between the low- and high-hierarchy regions is significantly correlated with 58 subject measures, which are much higher than the other three thickness measures. The dashed line denotes the average number of significantly correlated subject measures for the 66 individual brain parcels. (**C**) For each pair of thickness measures, a scatter plot shows their respective correlation strength (absolute correlation) with single subject measures that are significantly correlated with both (one circle per subject measure). The thickness ratio between the low- and high-hierarchy regions has a higher correlation with all the subject measures, as compared with the other three thickness measures.

It is worth noting that the above results presented information overlapped with the findings from the thickness-behavior CCA mode (**Fig. 1**), and was intended more for a quantitative comparison between the cross-hierarchy contrast and region-based measures of thickness in terms of their relationship with behavioral data. To avoid the circularity, we repeated the same analysis using the two groups of subjects from the split-half test (**Fig. S2**). The data from the first group was used as a test set for defining the lower- and higher-order regions using the CCA analysis, and the second half of the data was then used as a validation set to compare various thickness measures in terms of their correlations with subject measures. Similar results were obtained (**Fig. S3**).

### Co-modulations of functional connectivity with behavior show a cross-hierarchy contrast

Given the negative-to-positive behavioral change along the thickness-behavior CCA mode is similar to what has been observed for the connectivity-behavior CCA mode (12), we wanted to know whether the functional connectivity modulation along the connectivity-behavior CCA mode shares any similar features with the cortical thickness modulation along the thickness-behavior CCA mode. Towards this goal, we first applied CCA to the functional connectivity and behavioral data from 818 HCP subjects. Consistent with the previous study (12), we found a single highly significant the CCA mode (*r* = 0.77, *p* < 1.0×10^−5^, corrected) between the two modalities, which shows opposite correlations with positive and negative subject traits (**Fig. S4**). The top ones are largely overlapped with those of the thickness-behavior CCA mode (**Fig. 1C**).

We then examined the modulation of functional connectivity along this connectivity-behavior CCA mode. Briefly, we identified functional connections, among those of the 200 brain regions, that are significantly modulated along the connectivity-behavior CCA mode, i.e., showing significant correlation (*p* < 0.05, uncorrected) with the connectivity-behavior CCA mode. These connections are then multiplied by its sign to quantify how much their strength is upregulated (i.e., the positive connectivity becomes more positive or negative ones become more negative) or downregulated along the CCA mode direction. For each brain region, we counted the numbers of significantly upregulated and downregulated connections respectively and mapped the results onto the brain surface (Note: one brain region has both upregulated and downregulated connections). It should be noted that the mapping was done based on parcels rather than the ICA maps of the 200 regions (12), because the ICA maps can introduce a strong spatial pattern regardless of the connectivity changes (**Fig. S5**). The two resulting maps represent the degree to which the overall functional connectivity strength changes (increase or decrease) towards high-scoring subjects with more positive traits. A cross-hierarchy contrast is evident in both maps but in an opposite manner. The higher-order regions, particularly DMN, showed the largest degree of connectivity strength increase but the smallest degree of connectivity strength decrease, whereas the lower-order sensory/motor areas had an opposite pattern of modulation (**Fig. 3A** and **3B**). The cross-hierarchy pattern of the functional connectivity modulation is even clearer in the difference between the two maps (**Fig. 3C**).

**Fig. 3.**
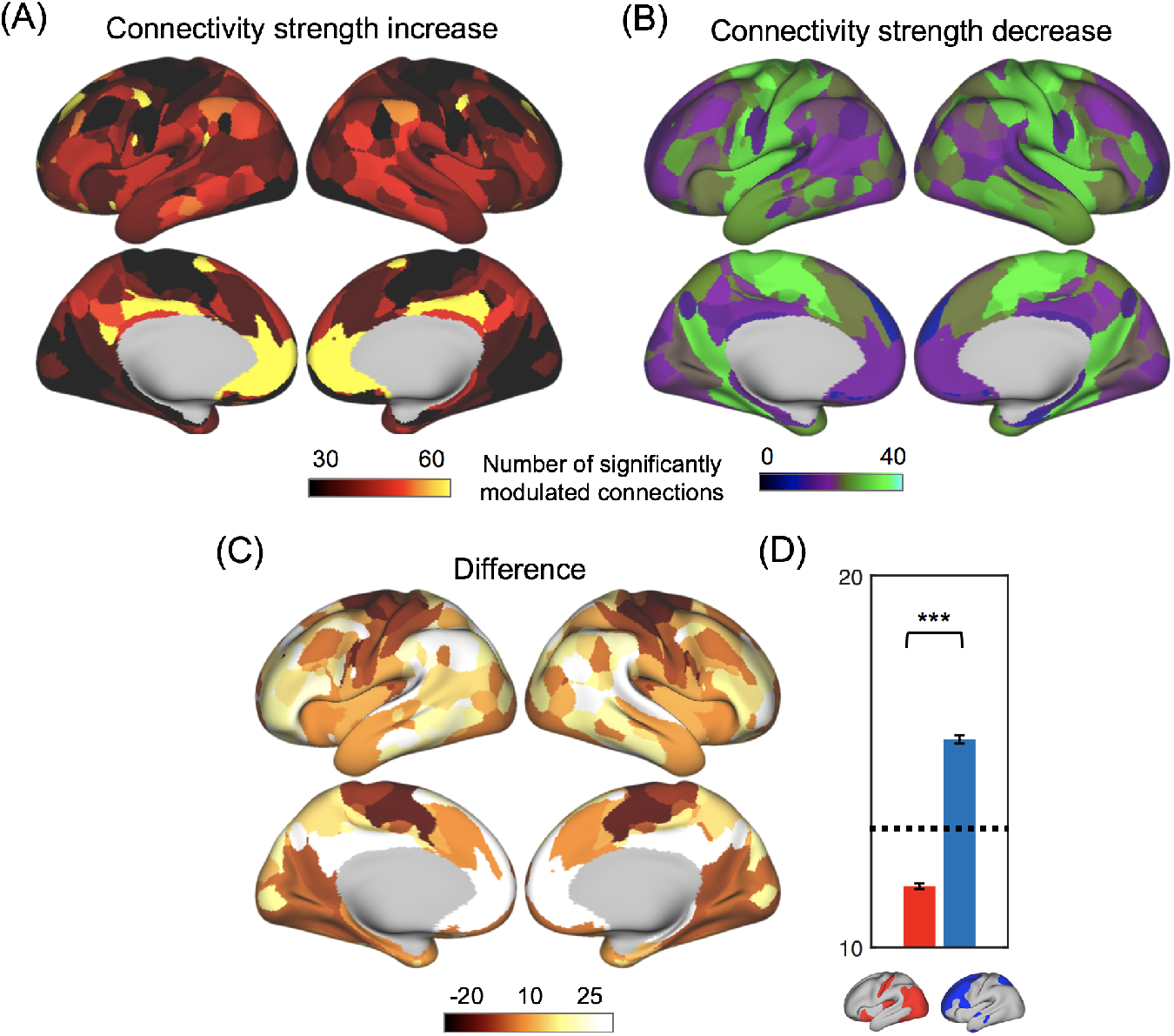
Functional connectivity strength modulations along the identified connectivity-behavior CCA mode. (**A**) Map of functional connectivity strength increase along the connectivity-behavior CCA mode towards subjects with more positive traits. The color of each brain region encodes the number of its connections (out of 199) whose strength increases significantly (i.e., upregulated) along the direction of the connectivity-behavior CCA mode. (**B**) Map of functional connectivity strength decrease along the connectivity-behavior CCA mode. The color of each brain region encodes the number of the connections whose strength decreases significantly (i.e., downregulated) along the direction of the connectivity-behavior CCA mode. (**C**) The difference map between the two shows how much more upregulated connections than downregulated ones for each region. (**D**) A bar plot summarizes the values of the difference map (**C**) within the lower-(blue) and high-order (red) brain regions. They are not only significantly different from each other (*p* = 0), but both are also significantly different (*p* = 0) from the mean values over the whole brain (the dashed line). Error bars represent the SEM across regions.

### A single direction featuring the maximal population covariation of the three modalities

The connectivity-behavior mode and thickness-behavior mode showed strong correlations with a similar set of subject measures (**Fig. 1C** and **Fig. S4**). In addition, the modulation of functional connectivity and cortical thickness along the two CCA modes displayed a similar divergent pattern across the cortical hierarchy (**Fig. 1D** and **Fig. 3C**). These correspondences suggest that the two modes may actually represent a similar direction in the subject space, which might also mediate the maximal correlation between the functional connectivity and cortical thickness. To have a complete understanding of the population covariation of the three modalities, we performed CCA between the functional connectivity and cortical thickness. Eight pairs of canonical variates survived through the statistical test (*p* < 0.05, corrected). The first pair with the highest correlation and the most significant *p*-value (*r* = 0.80, *p* <1.0×10^−5^, corrected) is associated with similar divergent modulations of both functional connectivity and cortical thickness (**Fig. 4**), suggesting it is closely related to the other two CCA modes. The 6 canonical variates from the three pairs of CCA show strong correlations with each other (correlation coefficients: 0.58 ± 0.13, all *p* < 7.5×10^−36^). We then performed PCA on the 6 canonical variates to obtain the first principal component to represent an eigendirection of the population covariation of the three modalities. This eigendirection accounts for 64.93% of the total variance of these 6 canonical variates and is highly correlated with each of them (correlation coefficients ranging from 0.78 to 0.82, all *p* ≤ 9.37×10^−170^) (**Fig. 4**). Overall, the findings suggest that the population variations of the cortical thickness, functional connectivity, and behavioral/demographic measures are maximally linked to each other along a similar direction in the subject space that is featured by an increase of overall goodness of subject measures and a divergent modulation of both functional connectivity and cortical thickness at the higher-order brain regions and low-level sensory/motor areas.

**Fig. 4.**
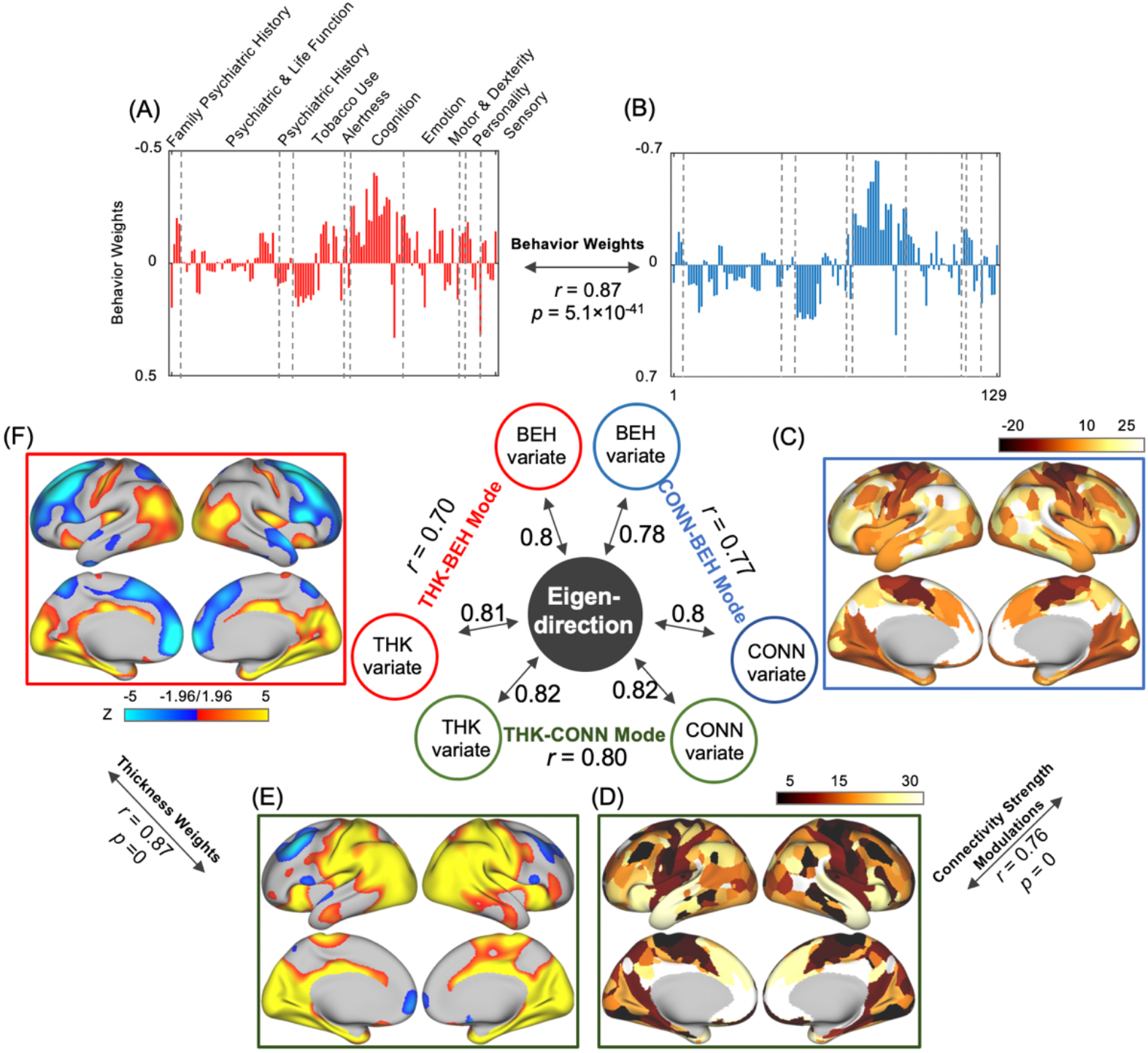
The relationships between the three pairwise CCA results. Three pairs of canonical variates from the three CCA show strong correlations with each other. The eigen-direction of the 6 canonical variates accounts for the majority of their variance (64.93%) and is strongly correlated with each one of them (correlation coefficients ranging from 0.78 to 0.82, all *p* ⩽ 9.37×10^−170^). The same modality of data, i.e., the behavioral/demographic measures (**A**-**B**), functional connectivity (**C**-**D**), and the cortical thickness (**E**-**F**), shows largely similar patterns of correlations with different CCA modes. All the correlations shown in this figure are statistically significant (all *p* < 1.0×10^−5^).

## Discussions

By applying a data-driven CCA to the cortical thickness, resting-state functional connectivity, and demographic/behavioral measures from a large cohort of subjects, we found that the maximal thickness-behavior correlation and the maximal connectivity-behavior correlation are converged in a specific direction, along which all three modalities show very characteristic modulations. Towards one end of this direction, individuals tend to have more positive subject traits as described previously (12), but more importantly, their functional connectivity and cortical thickness show a similar divergent modulation across the cortical hierarchy, i.e., increased functional connectivity and thinner cortex in the higher-order brain regions whereas decreased connectivity and thicker cortex at the low-level sensory/motor areas. Modern neuroimaging has been successful in linking specific aspects of brain function and/or human behavior to specific brain networks/regions. Based on the findings in this study, we argue that structural and functional contrast between the lower- and higher-order brain regions might be the brain feature tightly linked to a specific behavioral dimension that quantifies the overall goodness of human behavior and demographics.

The lower- and higher-order cortical regions showing opposite modulation in our CCA results are known to be structurally and functionally distinct. The higher-order brain regions expand much more as the brain size increases (26), over the developmental course (27), and also across species from monkeys to humans (27). In contrast, the lower-order sensory regions have a higher level of cortical differentiation and thus more refined layers (28). Cortical myelination follows a trajectory from the lower-order regions to the higher-order regions and thus creates a strong contrast between the two, which can be readily assessed by MRI-derived T1-weighted/T2-weighted maps (29). These structural differences, as well as other known differences in cytoarchitecture, cell types and synaptic physiology (30), between the two sets of brain regions are consistent with their distinct gene expression profiles (26, 30). Altogether, they may serve as the basis of their distinct functions, including those seen with neuroimaging tools. For example, embedding resting-state functional connectivity matrix into a low-dimensional space revealed a principal gradient separating the lower-order sensory/motor regions from the higher-order DMN (31). Meta-analysis of numerous neuroimaging literature identified a functional spectrum along this hierarchical gradient from simple sensory perceptions to abstract and complex cognitive functions (31). A series of studies have found consistent evidence across species that the resting-state connectivity/dynamics are divergently modulated across the lower- and higher-order regions from wake to anesthetized conditions (32–35). In addition, resting-state fMRI connectivity changes associated with a general psychopathological score, which reflects chronicity and symptom severity of mental illness in patient groups but risks in healthy individuals, also diverge in these two sets of brain regions (36).

The cortical thickness has been linked to various aspects of the human behavior, particularly the human intelligence (also seen in **Fig. 1A**), measured either with the intelligent quotient (IQ) or a general intelligence g-factor quantifying more general cognitive capabilities, by multiple studies (16–18, 37, 38). Most of them reported a positive association between the cortical thickness and intelligence. Although the spatial pattern of correlations varied across studies, relatively consistent correlations were observed in the lower-order sensory/motor regions (16, 18, 37, 39). The overall thickness-behavior correlations in the present study (**Fig. 1D**) biased slightly towards positive, which is somewhat consistent with the previous observations. However, significant negative correlations between the intelligence and the cortical thickness of higher-order brain regions were largely absent in the existing literature. There could be multiple reasons. First, the set of subject measures we used include real-life functions and demographic measures, such as education, income and life satisfaction, which can contribute significantly to the identified CCA modes. Indeed, the cortical thinning has been observed at the middle and superior frontal gyrus in subjects with high-level life satisfaction (20), as well as the primary sensory/motor areas in adults with psychopathy or violent antisocial personality disorder (21, 22) or school children showing aggressive behaviors (19). These findings are to a large extent in accordance with the divergent changes of the cortical thickness we found here. However, the contribution of the life-related factors cannot explain significant negative correlations between the thickness at some higher-order cortices, such as the frontal regions, and single measures of fluid intelligence or picture vocabulary (**Fig. S1C-E**). Thus, the discrepancy might also be due to other factors, such as differences in the cohort size, data quality, and pre-processing algorithms. In addition, a dynamic thickness-intelligence relationship across the lifespan and the relatively young age (28.8 ± 3.7 years old) of the HCP subjects might also contribute to this discrepancy (40).

Neural mechanisms underlying the divergent modulation of the cortical thickness are unclear, but there are a few possibilities based on previous literature. The cortical thinning is a process featuring the developmental course and has been hypothesized to reflect a neuronal pruning process of removing inefficient synapses (3, 38). In addition, intracortical myelination can also result in the apparent thinning of the MRI-reconstructed cortex (38). These microstructural changes might underlie the association between positive subject traits and the thinning of the higher-order cortex. In contrast, other region-specific processes may be involved in determining the thickness of the primary sensory/motor cortices, which showed an opposite modulation in the present study and much less significant change across the lifespan compared with the higher-order brain regions (38). As an example, the loss of mirror neurons has been proposed to be a possible reason for the thinner primary motor cortex associated with aggressive and violent behaviors (41). However, these explanations remain conjectures before proven.

The present study has a few limitations. First, this is a purely correlative study and the caution should be exercised when one attempts to infer any causal relationships based on the results. For example, the divergent modulations across the cortical hierarchy along the CCA mode direction were observed both for the cortical thickness and functional connectivity. A parsimonious interpretation would be that the structural difference leads to functional changes. However, we should not exclude the possibility that spontaneous brain activity underlying the resting-state functional connectivity may help to shape brain structure and morphology, such as the cortical thickness. Secondly, the divergent cortical thickness modulations at the lower- and higher-order brain regions might be, to some degree, disassociated in subpopulations. The cortical thickening at the lower-order sensory/motor regions is more prominent in subjects with more negative traits, whereas the cortical thinning at the higher-order regions appears to be stronger at the other end of the CCA mode (**Fig. 1E**). Lastly, we excluded behavioral measures of alcohol use in our CCA models, because we are not sure whether the alcohol use in the HCP subject cohort should be regarded as a positive or negative subject trait. Our preliminary analysis suggests that alcohol use is positively correlated with the CCA mode and other positive traits, such as fluid intelligence, which is against the common perception of it as a substance use. However, the inclusion of the measures of alcohol use produced little effects on our overall results, particularly the divergent pattern of cortical thickness modulations (**Fig. S6**).

In summary, the cortical thickness, functional connectivity, and behavioral/demographic measures from a large cohort of 818 HCP subjects show maximal correlations with each other along a similar direction across individuals, indicating a convergence of cross-modality relationships among these three types of data. This mode direction is associated with characteristic modulations in all three modalities, including a change in the overall goodness of behavioral and demographic measures and divergent modulations of both cortical thickness and functional connectivity across the cortical hierarchy between the lower- and higher-order brain regions. These findings suggest that the cross-hierarchy contrast of brain structural and functional properties may be an important feature tightly linked to a specific aspect, i.e., the overall goodness, of human behavior and demographics.

## Materials and Methods

### Data and preprocessing steps

We used demographic and behavioral measures, structural MRI and resting-state fMRI data from 818 HCP subjects. All subjects were healthy adults (28.8 ± 3.7 years old, 365 females). The demographic and behavioral measures include both “open access” and “restricted” items from the HCP dataset (http://humanconnectome.org/data). The structural and functional imaging data was preprocessed with the HCP minimal preprocessing pipelines (10) (see **SI Materials and Methods** for specific steps). We used the cortical thickness of the 818 subjects provided by the HCP as the main structural measure, which has been mapped onto the brain surface with 32K resolution, and performed additional spatial smoothing (FWHM = 6mm) on each thickness surface map. We obtained the resting-state functional connectivity of 200 distinct brain regions directly from the HCP PTN (Parcellation + Timeseries + Netmats) dataset (12, 42), which is publicly available (https://db.humanconnectome.org). Following the previous study (12), we used resting-state fMRI connectivity measures based on partial correlations of 200 ICA parcels for all the results presented in the paper.

### CCA between cortical thickness and behavioral/demographic measures

We used the CCA method (11) to identify the maximal linear correlation between cortical thickness at all surface vertices and 129 behavioral/demographic measures. Each significant CCA mode identifies a linear combination of thickness measurements and a linear combination of the 129 behavioral/demographic measures (see details in **SI Materials and Methods** for how this subset was selected), which are maximally correlated with each other across subjects. Eleven confounding factors, including age, gender, mean head motion, weight, height and blood pressure, and squared measures for 9 of these 11 factors (the other two were binary) were regressed out. The principal component analysis (PCA) was performed to reduce non-subject dimensions to 100, which has been used in the previous study (12), in order to avoid overfitting in CCA. Other principal component numbers, such as 60, 80 and 120, were also applied to validate the reproducibility of the above CCA (**Fig. S7**). The CCA was implemented by the Matlab function *canoncorr*. The principal CCA mode was identified as the first pair of canonical variates with the highest correlation. For statistical inference, we permuted subject ID for one modality of data (subject measures) by 100,000 times and run CCA for each permutation to build a null distribution. Multiple comparisons were corrected using the family-wise error rate (FWER), following the same procedure in the previous study (12). To examine changes in cortical thickness along the direction of the identified CCA mode, we defined a CCA score for the identified CCA mode as the mean of the corresponding pair of canonical variates. (see **SI Materials and Methods** for details)

To make sure that the overall relationship between the cortical thickness and behavior is not due to any single subject measures, we employed a leave-one-out procedure to re-run the CCA with excluding one of 129 subject measures each time (**Fig. S1**). To demonstrate the reproducibility of the thickness-behavior CCA and make sure that the overall results are not due to any specific subjects, the split-half test were applied, and we repeated our analysis on two randomly split samples (each with 409 subjects) and obtained the similar spatial patterns of thickness weights with that from 818 subjects (**Fig. S2**).

### CCA between functional connectivity and behavioral/demographic measures

We replicated the connectome-behavior CCA for 818 subjects using the same procedures described previously (12). We correlated the canonical variates with single connectivity and subject measures to obtain their canonical weights. We multiplied the connectivity weights by the sign of population mean connectivity across subjects to estimate the modulation of connectivity strength along the connectivity-behavior CCA mode. Then, we counted the number of significantly (*p* < 0.05, uncorrected) up- and down-modulated connections for each brain ROIs, and mapped the connectivity strength modulation along the CCA direction onto the brain surface (see details in **SI Materials and Methods**)

### CCA modeling of cortical thickness and functional connectivity

The same CCA procedures were also applied to the cortical thickness and functional connectivity from 200-dimensional group-ICA Netmat to derive thickness-connectivity CCA modes. Similar to the other two CCA models, we also calculated the correlations between the first CCA mode with the original sets of the cortical thickness and functional connectivity data to obtain their corresponding weights (**Fig. 4D** and **4E**).

### Eigen direction for the population covariance of three modalities

To understand the relationship among the principal CCA modes derived for the three pairs of different modalities of data, we performed PCA on the 6 canonical variates of the three CCA modes and took the first principal component (explains most total variance) as the eigen-direction of the population covariance of these three modalities of data. We correlated this eigen-direction with each of these 6 canonical variates. We further examined the correspondence among these CCA modes by computing the correlation between the weights of the same modality of data in different principal CCA modes, such as the behavioral weight of the thickness-behavior CCA mode and that of the connectome-behavior CCA mode respectively (**Fig. 4**).

More details are available in **SI Materials and Methods**.

## Acknowledgments

This research was supported by the NIH Pathway to Independence Award (K99/R00) 4R00NS092996-02.

## Author contributions

X.L., F.H., and Y.G. contributed to the data acquisition and image processing. F.H. and X.L. designed and carried out the statistical analyses. All authors contributed to writing the paper.

## Data and materials availability

The demographic, behavioral measures, cortical thickness and functional connectome data are from the HCP dataset (http://humanconnectome.org/data). All data is available in the main text or the supplementary materials.

