## supplemental table 1 for "Neuroimaging contrast across the cortical hierarchy is the feature maximally linked to behavior and demographics"

**List of Supplementary Materials**


Supplementary Materials and Methods
Figs. S1 to S7
Tables S1
References for supplementary materials

### Supplementary Materials and Methods

**Data and preprocessing steps.** We used structural MRI and resting-state fMRI data from 818 HCP subjects. All subjects were healthy adults (28.8 ± 3.7 years old, 365 females), who underwent one imaging sessions of structural MRI (T1w_MPR1 series, 3D MPRAGE, TE=2.14 ms, TR=2400 ms, 0.7 mm spatial resolution) and 4 sessions of resting-state fMRI (15 minutes each, TE=33.1ms, TR= 720 ms, multiband acceleration factor of 8) scanning on a 3T Siemens connectome-Skyra scanner. The demographic and behavioral measures include both "open access" and "restricted" items from the HCP dataset (http://humanconnectome.org/data).

The functional and structural imaging data was preprocessed with the HCP minimal preprocessing pipelines (1). Briefly, structured artifacts were identified and removed for each resting-state fMRI run by independent component analysis (ICA) followed by FMRIB's ICA based X-noiseifier (2, 3), which can remove 99% artifactual ICA components in each dataset. The resting-state fMRI data was mapped as time series of grayordinates, including cortical surface vertices and spatially standardized subcortical voxels. We obtained the resting-state functional connectivity of 200 distinct brain regions directly from the HCP PTN (Parcellation + Timeseries + Netmats) dataset (4), which is publicly available (https://db.humanconnectome.org). The dataset includes the resting-state connectivity measures for brain parcels ("nodes") derived with the ICA at several different dimensionalities (50, 100, 200, 300). Following the previous study (4), we used the resting-state fMRI connectivity measure based on partial correlations (5) of 200 ICA parcels for all the results presented in the paper, and we also tested the results from different ICA parcels described as below. Notably, the use of full Pearson's correlations did not change the major results and thus the conclusions of this paper. Refer to the previous report (4) about details of deriving the ICA parcellations and associated resting-state fMRI connectivity measures. The preprocessing of the structural imaging includes the T1w and T2w images alignment, bias field correction, the volume space registration into MNI space, white and pial cortical surface reconstruction, which derived the cortical thickness over the entire neocortex. We further performed spatial smoothing (FWHM = 6mm) on cortical thickness surface maps of 32K resolution.

**CCA between cortical thickness and behavioral/demographic measures.** We used the CCA method (6) to identify the maximal linear correlation between cortical thickness at all surface vertices and 129 behavioral/demographic measures. Each significant CCA mode identifies a linear combination of thickness measurements and a linear combination of behavioral/demographic measures, which are maximally correlated with each other across subjects. Specifically, thickness on the surface of 32K resolution (818 subjects × 59,412 vertices as matrix $T_1$) and 129 behavioral and demographic measures (from the original set of 478 measures, 818 subjects × 478 measures as $S_1$) were linked together using CCA.

We also generated an additional matrix $T_2$, where each column of $T_1$ was normalized according to its mean, and removed the badly conditioned columns with a low mean value (less than 0.1). Column-wise demeaning and matrix-wise variance-normalization were operated for $T_1$ and $T_2$, respectively, and then those two resulting matrices were concatenated horizontally to give rise to a matrix $T_3$ that includes both preferences as the previous study (4). An inverse Gaussian transformation based on ranking was used to promote Gaussianity of $S_1$ and generate $S_2$, by which the effect from potential outliers would be decreased.

11 behavioral/demographic measures were considered as the confound factors:

1. Acquisition reconstruction software version.
2. Gender.
3. Age.
4. Mean head motion (7).
5. Weight.
6. Height.
7. Systolic blood pressure.
8. Diastolic blood pressure.
9. Blood hemoglobin A1C measures.
10. The cube-root of brain volume (ventricles included, estimated by FreeSurfer).
11. The cube-root of intracranial volume (estimated by FreeSurfer).

Besides those 11 measures, we demeaned and squared the measures 3-11 (since the first two are binary), and then took them as another set of confounding factors so that we can remove the potentially nonlinear effects of those measures. All 20 confound factors were demeaned and then regressed out from the $T_3$and $S_2$.

Race was excluded from the analysis similar to the previous study (4). Several subject measures were also excluded before CCA as described:

1. The 11 confound behavior measures listed above.
2. 88 poorly quantified measures according to the following criteria (4):
    a. Fewer than 250 subjects with valid measures;
    b. Measures that >95% of subjects had the identical scores;
    c. The measures had extreme outlier value. For $x_s$ as the score of subject $s$ for a subject measure, we calculated $y_s = (x_s - \text{median}(x_s))^2$. If $\max(y_s) > 100 \times \text{mean}(y_s)$ , we regarded the measure as having extreme outliers.
3. 45 variables that are perceived irrelevant to brain function (For example, "Is the subject born in Missouri?" or "Is the subject in a live-in relationship?") or highly dependent variables (i.e. Body mass index is dependent on height and weight). All of those 45 variables are same to a previous study (4).
4. 191 imaging measures that are not behavioral traits (i.e. cortical thickness, regional volume and surface area).
5. 22 measures related to the alcohol use. Although the inclusion of alcohol measures produced very similar CCA results (**Fig. S6**), the alcohol usage exhibited similar correlations as positive personal traits with the cortical thickness and functional connectivity. Although the finding is consistent with their positive correlations with fluid intelligence, which were found in both the current study and previous ones (8), it is somewhat against the common perception of alcohol usage as a substance use. To avoid the complications, we exclude the measures related to alcohol use from the main analysis.

The above procedures gave rise to a 818 subjects × 129 subject measures matrix $S_3$. We list those 129 measures in **Table S1**. Importantly, the principal component analysis (PCA) was performed to reduce non-subject dimensions to 100, which is inspired by the previous study (4), in order to avoid overfitting in CCA. That is, the first 100 principal components of both subject measures and thickness (named $T_4$ and $S_4$, respectively, both were 818×100 in size) were used for thickness-behavior CCA. Notably, the $T_4$ and $S_4$ accounted for the 99.72% and 82.28% of the total variance of behavior and thickness data respectively. Notably, other principal component numbers, such as 60, 80 and 120, were also applied to validate the reproducibility of the above CCA (**Fig. S7**).

The CCA, implemented by the Matlab function *canoncorr*, estimated 100 modes by optimizing the de-mixing matrices $A_{TS}$ and $B_{TS}$ to obtain the maximal similarity between the two matrices $U_{TS} = T_4 A_{TS}$ and $V_{TS} = S_4 B_{TS}$ (both were $818 \times 100$ in size), and each pair of corresponding columns from $U_{TS}$ and $V_{TS}$ represents the strength with which a mode of population variation is common to subject measures and thickness. For statistical inference, we permuted the index of subjects (the rows of $T_4$ or $S_4$) by 100,000 times without removing the family structure as previous study (4) (since the 818 HCP subjects are from hundreds of families) and run CCA for each permutation to build a null distribution (**Fig. 1B**).

The principal CCA mode was identified as the first pair of canonical variates with the highest correlation ($r = 0.70$, $p < 1.0 \times 10^{-5}$, corrected for multiple comparisons across all modes estimated), which was the only significant mode ($p < 0.05$) among all 100 modes. Multiple comparisons were corrected using the family-wise error rate (FWER) referred to the Matlab script provided by previous study (4). The canonical variates of this principal mode, i.e., the first columns of $U_{TS}$ ($818 \times 1$) and $V_{TS}$ ($818 \times 1$), were correlated with cortical thickness ($818 \times 59{,}412$) and subject measures ($818 \times 478$), respectively, and the resulting weights indicate the contributions of the cortical thickness at single vertices and individual subject measures to the identified CCA mode. The map of thickness correlations (thickness weights, $59{,}412 \times 1$) was further converted to Z score using Fisher's transformation.

To examine changes in cortical thickness along the direction of the identified CCA mode, we defined a CCA score for the identified CCA mode as the mean of the corresponding pair of canonical variates, i.e., the first columns of $U_{TS}$ and $V_{TS}$. The 818 subjects were divided into four groups according to the quartiles of their CCA scores. We also defined two masks covering the lower (positively correlated with the principal CCA, $p < 0.05$, uncorrected, warm color portion in **Fig. 1D**) and higher (negatively correlated with the principal CCA mode, $p < 0.05$, uncorrected, cold color portion in **Fig. 1D**) hierarchical regions, respectively. The cortical thickness was averaged first within these two masks and then across subjects within each of the four groups to show the divergent modulations of cortical thickness across the cortical hierarchy along the identified CCA mode (**Fig. 1E**).

To make sure that the overall relationship between the thickness and behavior is not due to any single subject measures, we employed a leave-one-out procedure to re-run the CCA with excluding one of 129 subject measures each time. We also derived the cortical thickness weights ($59{,}412 \times 1$) for each reduced principal CCA mode and then correlated these 129 thickness weights results ($59{,}412 \times 129$) with thickness weights derived from the original one without leave-one-out ($59{,}412 \times 1$) (**Fig. S1B**). In addition, we also directly correlated 8 leading subject measures ($818 \times 8$, 8 subject measures showing the strongest correlations (absolute value) in **Fig. 1C**) with the cortical thickness measures ($818 \times 59{,}412$) across 818 subjects (**Fig. S1C-J**) to derive the thickness weights for each of these 8 subject measures.

We demonstrated the reproducibility of the thickness-behavior CCA and confirmed that the results are not from any specific subject with a split-half test. To be specific, we replicated our analysis on two randomly split samples (each was in size of 409 subjects) and then obtained the spatial patterns of (Fisher) z-transformed thickness weights for both first and second half group shown as **Fig. S2**.

To test whether the cross-hierarchy contrast of the cortical thickness is more closely related with the subject measures than region-based measures, we calculated the mean cortical thickness within the low- and high-hierarchy masks shown in **Fig. 1D** and **1E**, as well as their ratio, and then correlated them (818×3) with 129 subject measures across all 818 subjects (818×129). The resulting correlations were sorted and compared with those derived using the mean cortical thickness of 66 pre-defined brain regions (9). To build a control case for the ratio index, we rotated the two masks on the spherical brain surface by randomly generated angles ([294, 327, 46] with respect to the x, y and z axis respectively). We then calculated the mean cortical thickness within the control masks and their ratio, and correlated the thickness ratio between the control masks with the behavioral/demographic measures. Each group of correlations results (from the 5 sets) was sorted based on their correlation coefficients (**Fig. 2A**). We then counted the number of significantly correlated ($p$ < 0.05, uncorrected) subject measures for each of four different thickness measures, including the mean thickness of the low-hierarchy mask, of the high-hierarchy mask, the thickness ratio between the low- and high-hierarchy masks, and between the two control masks (**Fig. 2B**). For subject measures showing significant correlations with more than one thickness measure, we compared their absolute correlations with a pair of thickness measures using scatter plots (**Fig. 2C**). To avoid circuity, we defined new lower- and higher-order masks based on the first half results of thickness weights and replicated the above analysis by comparing various thickness measures (within the new masks) in terms of their correlations with subject measures (**Fig. S3**).

**CCA between functional connectivity and behavioral/demographic measures.** We replicated the connectome-behavior CCA (4) for 818 subjects. Functional connectivity data was obtained from PTN dataset described above. To be specific, the Netmat from 200-dimensional group-ICA was used as the connectivity matrix. The 200×200 Netmat matrices give functional connectivity (partial correlations converted to Z scores already) for 19,900 pairs (upper triangular matrix without diagonals) of brain regions. Matrices of functional connectivity ($C_1$, 818×19,900) and subject measures ($S_1$, 818×129) were constructed for all the subjects. Using the same procedures described above, we performed CCA and obtained a highly significant connectome-behavior CCA mode ($r$ = 0.77, $p < 1.0\times10^{-5}$, corrected for multiple comparisons). We also correlated the corresponding canonical variates in this significant (principal) CCA mode, i.e., the first columns of $U_{CS1}$ and $V_{CS1}$, with the original sets of connectivity $C_1$ and subject measures $S_1$ respectively and obtained the connectivity weights $A_{CS}$ and behavior weights $B_{CS}$ to show how the connectome-behavior CCA mode is correlated with functional connection between each pairwise brain regions and single subject measure. Since the connectivity metric can be of negative values, its negative correlation with the CCA canonical variate may actually indicate as a decrease of connectivity strength along the direction of CCA mode. We thus multiplied each connectivity weight, i.e., the correlation between single connectivity and the CCA canonical variate, by the sign of population mean connectivity (averaged $C_1$ across each column), and the results quantify how the strength (i.e., amplitude) of single functional connectivity is correlated with the identified CCA mode. Then, we mapped the connectivity strength modulations (along the CCA direction) onto the brain surface by counting, for each brain parcel, the number of connections (out of 199 in total) showing significantly positive (increased connectivity strength with increased CCA scores) and negative (decreased connectivity strength with increased CCA scores) correlations with the CCA connectivity canonical variate (**Fig. 3A** and **3B**). The resulting "connectivity strength decrease" map was also subtracted from the "connectivity strength increase" map to derive a "difference" map for functional connectivity modulations (**Fig. 3C**), whose map scores were averaged within the higher- and lower-hierarchy masks respectively. This parcel-based mapping method is modified based on the previous one, which multiplied the functional connectivity strength modulation of each ROI with its spatial ICA map and then summed them up to obtain a final map. Although the previous approach gives a smooth transition across the cortex, we found that the resulting maps all share a spatial pattern with the simple summation of all the ICA maps (**Fig. S5**).

To increase confidence that the connectome-behavior CCA mode is robust, we replicated the CCA on the same subject measures and Netmat (connectivity) derived from 50-, 100- and 300-dimensional group-ICA parcellation respectively. Similar to the previous study (4), we examined the following correlations between the original mode (200-dimensional ICA parcellation) and the alternative one:

1. The connectivity variates in principal CCA mode (the most significant one with maximal correlation): $Ucs_1$(original) versus $Ucs_1$(alternative);
2. The behavior variates in principal CCA mode: $Vcs_1$(original) versus $Vcs_1$(alternative);
3. The behavior weights derived by the principal CCA mode: *Bcs*(original) versus *Bcs*(alternative);

The connectivity weights *Acs* was excluded in the comparison due to incompatible dimensions between the original and alternative one. We thus obtained the above three correlations between the original mode and the 50-dimensional one as [0.56 0.75 0.88], the 100-dimensional one as [0.63 0.82 0.93] or the 300-dimensional one as [0.60 0.81 0.94] (all $p$-values $< 2.45\times10^{-67}$, uncorrected).

**CCA modeling of cortical thickness and functional connectivity.** Using the same procedures as the other two CCA modeling, we applied the CCA to the thickness data and Netmat from 200-dimensional group-ICA (200×200) to derive thickness-connectivity CCA modes. This produced 8 significant CCA modes in which the first one had the highest correlation ($r = 0.80$, $p < 1.0\times10^{-5}$). We also correlated local cortical thickness and pairwise functional connections to the first CCA mode, i.e., the corresponding canonical variates, to obtain the canonical weights for both modalities (**Fig. 4 D** and **4E**).

**Eigen direction for the population covariance of three modalities.** We performed PCA on the 6 canonical variates from the three CCA modes and took the first principal component (explains 64.93% of the total variance) as an eigen-direction of population covariance of these three modalities of data. We validated this eigen-direction by correlating it with these 6 canonical variates from three principal CCA modes. We also computed the correlation between the two behavioral weights from principal thickness-behavior CCA mode and principal connectome-behavior CCA mode, the correlation between the two thickness weights from principal thickness-behavior CCA mode and principal thickness-connectome CCA mode, and the correlation between the two connectome weights from principal connectome-behavior CCA mode and principal thickness-connectome CCA mode (**Fig. 4**).

**Supplementary Figures**

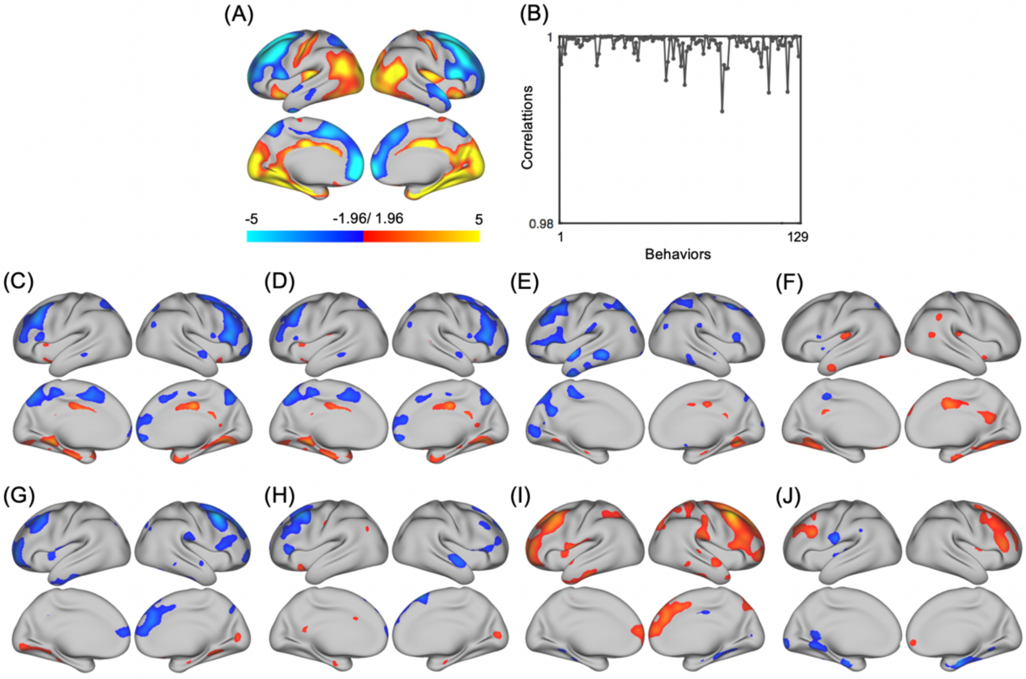


**Fig. S1. The contribution of single subject measures on the thickness-behavior CCA mode.** (**A**) The brain surface map of significant Fisher-z transformed thickness weights (correlations between local thickness and identified CCA mode, $p < 0.05$, uncorrected, same as **Fig. 1D**). (**B**) Spatial similarity between the original thickness weights map (**A**) with those obtained with leaving one subject measure out of analysis. There are almost no changes as evidenced by spatial correlations close to 1. (**C**-**J**) Spatial maps showing correlations (Fisher-z transformed thickness weights) between the cortical thickness and single subject measures. The maps are for the top 8 subject measures that show the strongest correlations with the thickness-behavior CCA mode, including **C**: Picture vocabulary test; **D**: Picture vocabulary test (age normed); **E**: Fluid intelligence: correct response in PMAT; **F**: Delay discounting (AUC for discounting of $40K); **G**: Cognition: PLO test total correct number; **H**: Picture sequence memory test (age normed); **I**: Cognition: PLO test total position off number; **J**: Audition: words in noise.

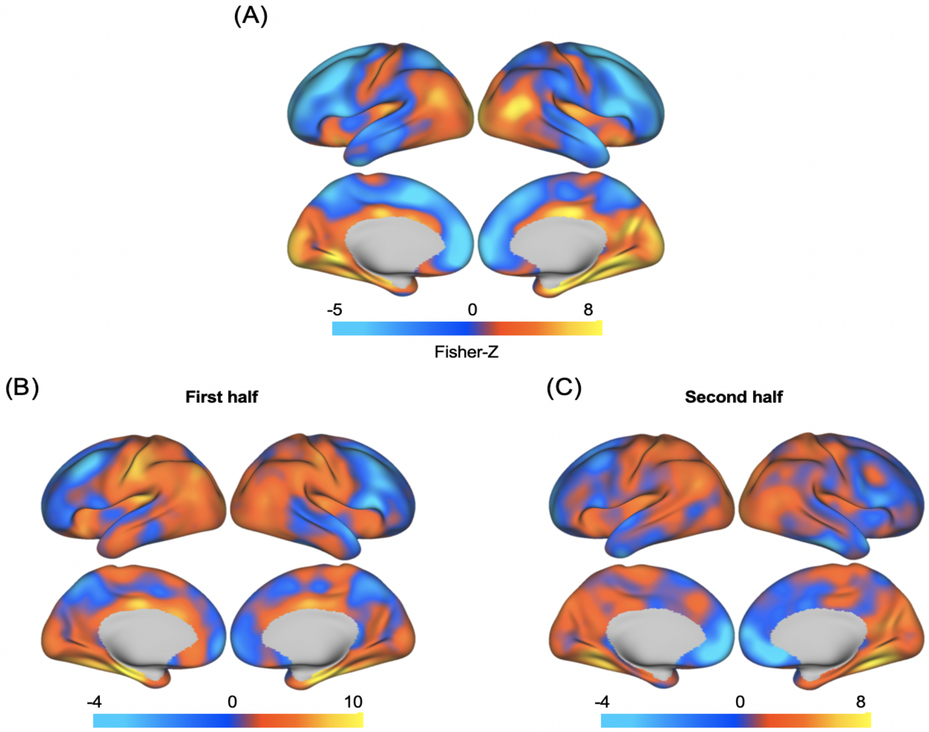


**Fig. S2. Split-half test for the thickness-behavior CCA mode.** To demonstrate the reproducibility of the thickness-behavior CCA, we randomly divided the 818 sample into the first half and second half, and derive thickness weights as panel **B** & **C** by following the method to derive **Fig. 1D** (as **A** in this figure). The original correlations are converted to Fisher's z scores. The correlation between map **A** and **B** (or **C**) across vertices were 0.752 and 0.704, respectively (all *p*-values were 0).

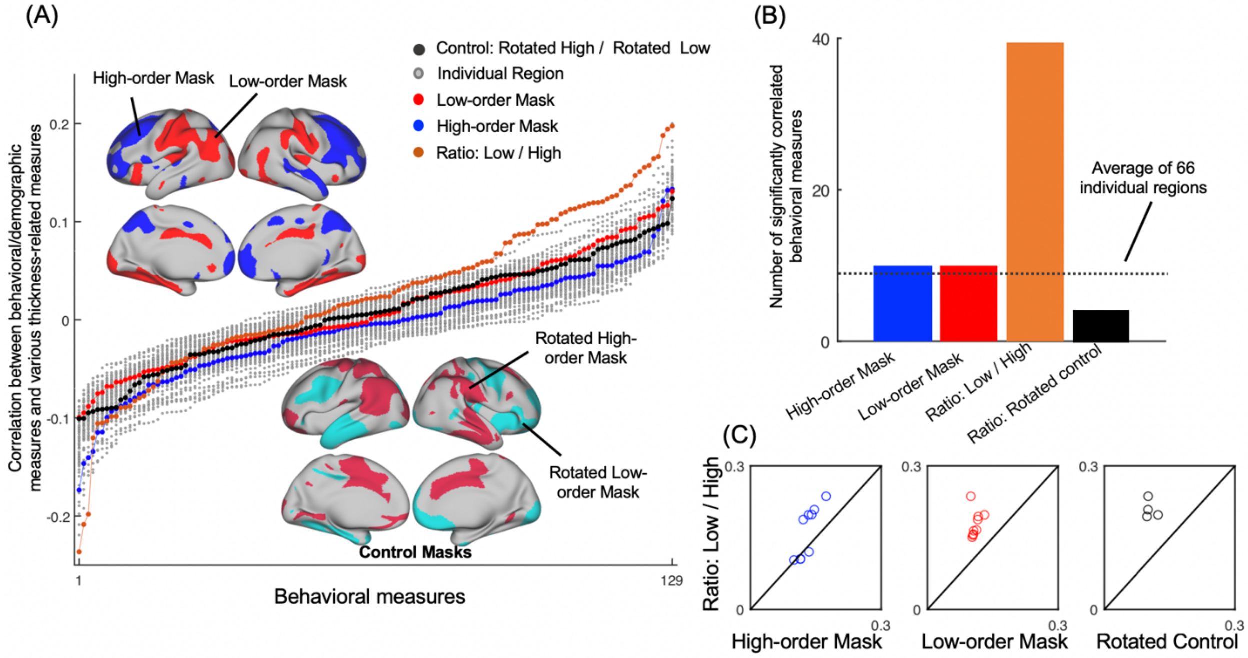


**Fig. S3. Correlations between subject measures and various thickness measures based on two halves of the data.** (**A**) Sorted correlations between 129 subject measures and 70 different thickness measures across the validation data (the second half subjects), including the cortical thickness of 66 pre-defined brain parcels (9) (gray lines), of the lower- (red line) and higher-order (blue line) masks, which were derived from the training data (the first half subjects from the half-half test), the thickness ratio between these two masks (orange line), as well as the thickness ratio between two control masks obtained by randomly rotating the lower- and higher-hierarchy masks on the brain surface (black line). (**B**) The number of significantly correlated subject measures ($p <$ 0.05, uncorrected) was counted for four thickness measures. The thickness ratio between the low- and high-hierarchy regions is significantly correlated with 39 subject measures, which are much higher than the other three thickness measures. The dashed line denotes the average number of significantly correlated subject measures for the 66 individual brain parcels. (**C**) For each pair of thickness measures, a scatter plot shows their respective correlation strength (absolute correlation) with single subject measures that are significantly correlated with both (one circle per subject measure). The thickness ratio between the low- and high-hierarchy regions has a higher correlation with all the subject measures, than the other three thickness measures.

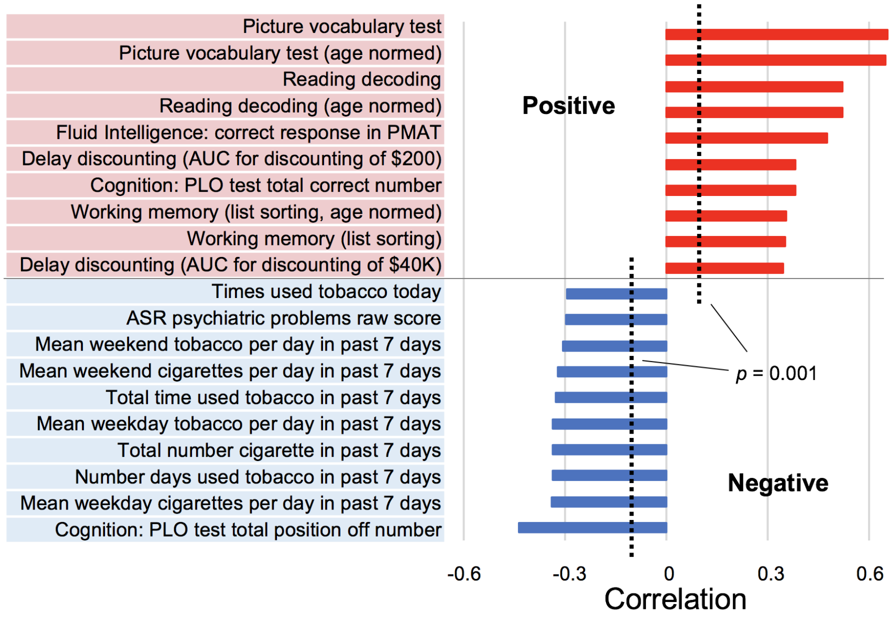


**Fig. S4. The top subject measures most strongly correlated with the identified connectome-behavior CCA mode.** The top positively correlated subject measures (red) generally describe positive personal traits, whereas the top negatively correlated subject measures (blue) are all associated with negative personal traits. Basically, this figure is demonstrated to validate the pattern that those positive traits are positively correlated with the identified connectome-behavior CCA mode, whereas negative traits are negatively correlated with the mode. Notably, although different naming way for the behavior or trait items with the previous study (4), most of the positive traits, the “PLO cognition test position off number” and tobacco use related items above are aligned with the results from previous one (4).

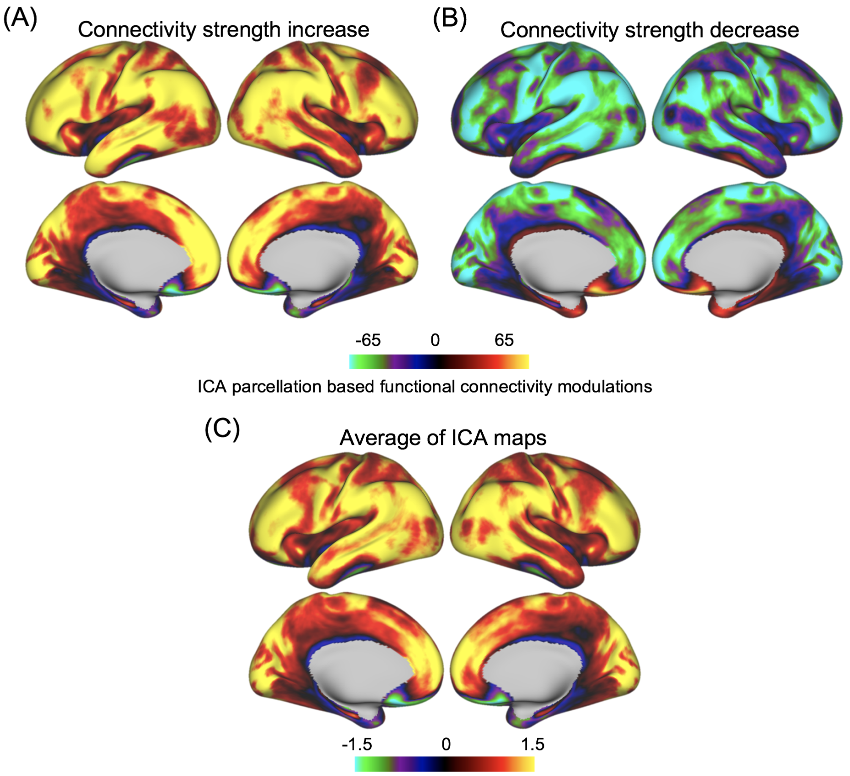


**Fig. S5. Mapping the functional connectivity strength modulations using the ICA maps of the brain parcels.** The parcel-based maps shown in **Fig. 3A** and **3B** were converted to **A** and **B** by using parcel-specific values as weights for averaging their corresponding ICA maps. While this method gives a spatially smooth map, the resulting patterns are largely dominated by the simple average of all the ICA maps without any weighting (**C**).

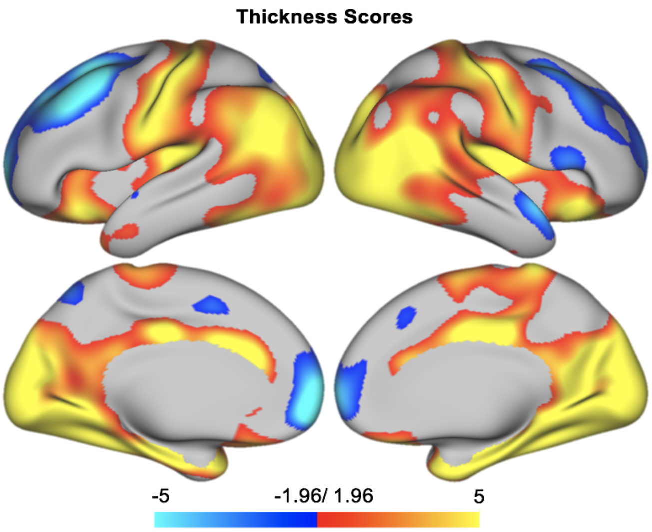


**Fig. S6. The thickness-behavior CCA mode with including the 22 measures related to alcohol use.** The way to derive this map is identical to **Fig. 1D** but includes the additional 22 subject measures related to alcohol use. The original correlations are converted to Fisher's z scores. Positive correlations (red-yellow colors) are mostly seen at the lower-level sensory/motor regions, whereas negative correlations (blue-cyan colors) are more dominant at the higher-order cognitive brain regions.

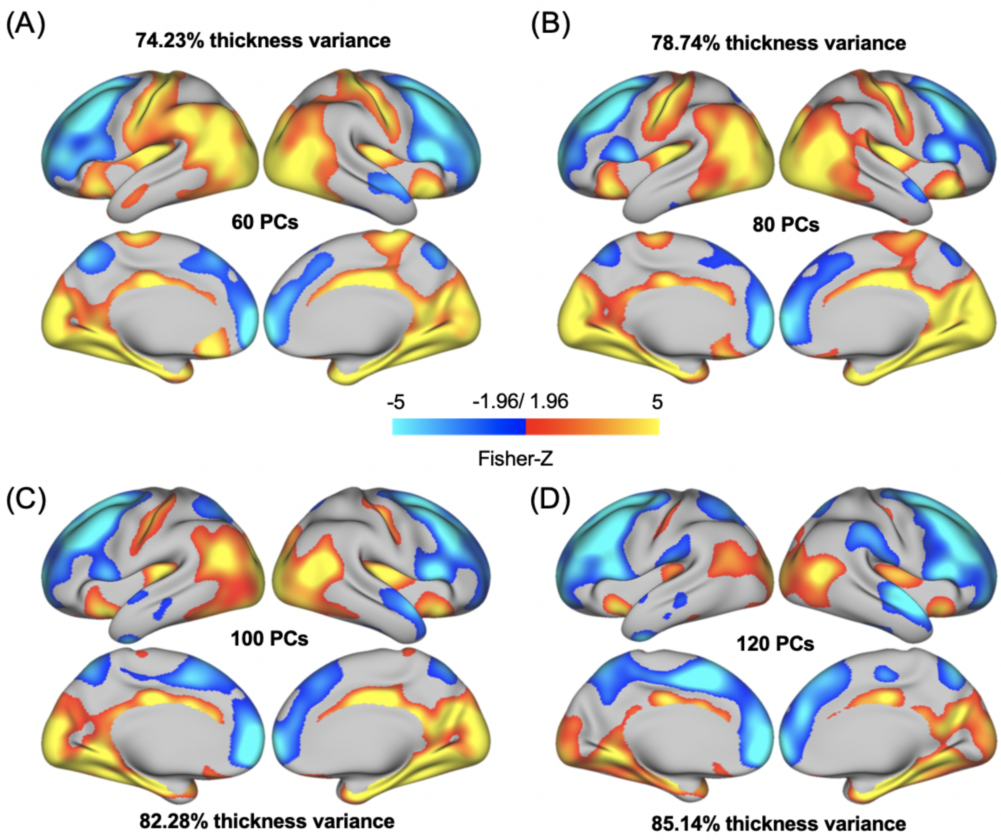


**Fig. S7. The thickness-behavior CCA modes with retaining different amount of data variance.** In addition to the 100 PCs retained for our original thickness-behavior CCA, the 60, 80 and 120 PCs for both thickness and subject measures variates in the PCA were used (respectively accounting for 74.23%, 78.74% and 85.14% of the total thickness variance, and 92.66%, 97.93% and 100% of the total subject measures variance) to examine the robustness of the PC numbers. The way to derive the figure was similar to **Fig. 1D** (100 PCs as **C**). The original correlations were converted to Fisher's z scores. The correlation between map **C** and **A** (or **B** or **D**) the across vertices were 0.927, 0.966 and 0.968, respectively (all *p*-values were 0).

**Table S1.** The list of 129 subject measures or demographics involved in CCA modeling (Alcohol use items excluded)

| **129 subject measures or demographics in CCA** | | |
|---|---|---|
| No.1-No.43 | No.44-No.86 | No.87-No.129 |
| 'FamHist_Moth_Dep' | 'SSAGA_ChildhoodConduct' | 'VSPLOT_TC' |
| 'FamHist_Fath_Dep' | 'SSAGA_PanicDisorder' | 'VSPLOT_CRTE' |
| 'FamHist_Moth_None' | 'SSAGA_Agoraphobia' | 'VSPLOT_OFF' |
| 'FamHist_Fath_None' | 'SSAGA_Depressive_Ep' | 'SCPT_SPEC' |
| 'ASR_Anxd_Raw' | 'SSAGA_Depressive_Sx' | 'IWRD_TOT' |
| 'ASR_Anxd_Pct' | 'Correction' | 'ListSort_Unadj' |
| 'ASR_Witd_Raw' | 'Total_Any_Tobacco_7days' | 'ListSort_AgeAdj' |
| 'ASR_Witd_Pct' | 'Times_Used_Any_Tobacco_Today' | 'ER40_CR' |
| 'ASR_Soma_Raw' | 'Num_Days_Used_Any_Tobacco_7days' | 'ER40ANG' |
| 'ASR_Soma_Pct' | 'Avg_Weekday_Any_Tobacco_7days' | 'ER40FEAR' |
| 'ASR_Thot_Raw' | 'Avg_Weekend_Any_Tobacco_7days' | 'ER40NOE' |
| 'ASR_Thot_Pct' | 'Total_Cigarettes_7days' | 'ER40SAD' |
| 'ASR_Attn_Raw' | 'Avg_Weekday_Cigarettes_7days' | 'AngAffect_Unadj' |
| 'ASR_Attn_Pct' | 'Avg_Weekend_Cigarettes_7days' | 'AngHostil_Unadj' |
| 'ASR_Aggr_Raw' | 'SSAGA_TB_Smoking_History' | 'AngAggr_Unadj' |
| 'ASR_Aggr_Pct' | 'SSAGA_TB_Still_Smoking' | 'FearAffect_Unadj' |
| 'ASR_Rule_Raw' | 'SSAGA_Times_Used_Illicits' | 'FearSomat_Unadj' |
| 'ASR_Rule_Pct' | 'SSAGA_Times_Used_Cocaine' | 'Sadness_Unadj' |
| 'ASR_Intr_Raw' | 'SSAGA_Times_Used_Hallucinogens' | 'LifeSatisf_Unadj' |
| 'ASR_Intr_Pct' | 'SSAGA_Times_Used_Opiates' | 'MeanPurp_Unadj' |
| 'ASR_Oth_Raw' | 'SSAGA_Times_Used_Sedatives' | 'PosAffect_Unadj' |
| 'ASR_Crit_Raw' | 'SSAGA_Times_Used_Stimulants' | 'Friendship_Unadj' |
| 'ASR_Intn_Raw' | 'SSAGA_Mj_Use' | 'Loneliness_Unadj' |
| 'ASR_Intn_T' | 'SSAGA_Mj_Ab_Dep' | 'PercHostil_Unadj' |
| 'ASR_Extn_Raw' | 'SSAGA_Mj_Age_1st_Use' | 'PercReject_Unadj' |
| 'ASR_Extn_T' | 'SSAGA_Mj_Times_Used' | 'EmotSupp_Unadj' |
| 'ASR_TAO_Sum' | 'MMSE_Score' | 'InstruSupp_Unadj' |
| 'ASR_Totp_Raw' | 'PSQI_Score' | 'PercStress_Unadj' |
| 'ASR_Totp_T' | 'PicSeq_Unadj' | 'SelfEff_Unadj' |
| 'DSM_Depr_Raw' | 'PicSeq_AgeAdj' | 'Dexterity_Unadj' |
| 'DSM_Depr_Pct' | 'CardSort_Unadj' | 'Dexterity_AgeAdj' |
| 'DSM_Anxi_Raw' | 'CardSort_AgeAdj' | 'NEOFAC_A' |
| 'DSM_Anxi_Pct' | 'Flanker_Unadj' | 'NEOFAC_O' |
| 'DSM_Somp_Raw' | 'Flanker_AgeAdj' | 'NEOFAC_C' |
| 'DSM_Somp_Pct' | 'PMAT24_A_CR' | 'NEOFAC_N' |
| 'DSM_Avoid_Raw' | 'ReadEng_Unadj' | 'NEOFAC_E' |
| 'DSM_Avoid_Pct' | 'ReadEng_AgeAdj' | 'Noise_Comp' |
| 'DSM_Adh_Raw' | 'PicVocab_Unadj' | 'Odor_Unadj' |
| 'DSM_Adh_Pct' | 'PicVocab_AgeAdj' | 'Odor_AgeAdj' |
| 'DSM_Inat_Raw' | 'ProcSpeed_Unadj' | 'PainInterf_Tscore' |
| 'DSM_Hype_Raw' | 'ProcSpeed_AgeAdj' | 'Taste_Unadj' |
| 'DSM_Antis_Raw' | 'DDisc_AUC_200' | 'Taste_AgeAdj' |
| 'DSM_Antis_Pct' | 'DDisc_AUC_40K' | 'Mars_Log_Score' |

**Note:** The full name of those items can be referred to the HCP 900 Subjects Release Reference Manual (https://www.humanconnectome.org/storage/app/media/documentation/s900/HCP_S900_Release_Reference_Manual.pdf) and HCP Data Dictionary (https://wiki.humanconnectome.org/display/PublicData/HCP+Data+Dictionary+Public-+Updated+for+the+1200+Subject+Release).
